# Early resource scarcity drives persistent transcriptional changes and vascular remodeling in the female prefrontal cortex

**DOI:** 10.64898/2026.06.11.731696

**Authors:** E. Andrews, R.C. Crist, A.I. Silva, S.N. Chehimi, G. Miranda, J. Shuey, L. Berhane, L. Hoot, E. Harris, A. Cuarenta, M.E. Wimmer, D. Bangasser, B.C. Reiner

## Abstract

Early childhood poverty is an environmental risk factor for psychiatric and neurodegenerative disorders, yet the cellular mechanisms by which resource scarcity produces persistent brain vulnerability remain poorly understood. The medial prefrontal cortex (mPFC), which regulates executive function and motivated behavior, is sensitive to early environmental conditions. To identify mechanisms linking early resource scarcity to lasting mPFC dysfunction, we used the rat limited bedding and nesting (LBN) model, which recapitulates key features of poverty. Prior work shows LBN disrupts mPFC-mediated behaviors in adulthood, often in a sex-specific manner. Here, we used single-nucleus RNA sequencing (snRNAseq) to identify sex- and cell-type-specific transcriptional alterations in the adult mPFC following brief postnatal LBN exposure or control housing. LBN induced more differentially expressed genes (DEGs) across multiple pyramidal neuron clusters in females than in males. Unexpectedly, the largest transcriptional changes due to LBN occurred in vascular cells in females, whereas male vascular cells exhibited no DEGs. These female-specific vascular genes were enriched for alterations of transcriptional programs regulating angiogenesis and endothelial structure. This molecular profile was orthogonally validated with 3D vascular reconstruction, revealing LBN reduced vascular coverage in the adult female mPFC, driven by decreased vessel volume and shortened vessel length, while males were unaffected. Reduced vascular coverage may constrain metabolic support to this region. The postnatal period is a critical window for vascular maturation, and, taken together, these findings identify persistent, female-specific vascular alterations as a novel and previously unrecognized mechanism through which early resource scarcity may persistently affect brain function and vulnerability.

## Introduction

Early environmental influences, such as poverty, parental care, sensory stimulation, nutrition, etc., can affect the risk of developing psychiatric and neurodegenerative disorders^1^. This relationship is mediated by the developmental reprogramming of the brain by stimuli in the early environment^2^. Given that brain regions mature at different rates, regions with extended developmental windows are particularly sensitive to postnatal environmental inputs. The medial prefrontal cortex (mPFC) exhibits protracted maturation into early adulthood and plays a key role in higher-order cognitive functions, motivational drive, and affective regulation^3,4^. These domains are disrupted in several neuropsychiatric and neurodegenerative disorders, including major depressive disorder, substance use disorder, and dementias. Thus, investigating how postnatal environmental stimuli influence the mPFC provides critical insights into the neurobiological mechanisms underlying risk for psychopathology.

In the human mPFC, research into how early adverse environmental exposures affect molecular signatures is constrained by ethical concerns about inducing early-life stress and by limited options for assessing molecular changes in the human brain^2^. Thus, non-human animal models have been developed. One widely used example is the limited bedding and nesting (LBN) manipulation, which simulates aspects of human poverty in rodents by restricting resources available to the dam and pups^5–7^. LBN also induces unpredictable maternal care, which is commonly observed in people parenting in under-resourced environments^5,8^. Prior rearing in LBN affects cognitive processes in adulthood that rely on the mPFC, including reversal learning, impulsivity, and risky decision-making^9–11^. Compared to rats raised with adequate resources, LBN rats also have altered motivated behaviors, such as juvenile play and adult motivation for social, sucrose, and drug rewards^10,12–16^ ; behaviors that engage the mPFC.

The lasting effects of LBN on adult behaviors are thought to be mediated by persistent transcriptional changes in key brain regions. Indeed, using bulk RNA sequencing (RNAseq), genes involved in glutamate signaling, immune signaling (e.g, IL-1, IL-6, IL-9) and intracellular signaling pathways (e.g., cAMP, phsophatidylionsitol) are altered in adult LBN vs. control rats in regions including the orbitofrontal cortex (OFC), nucleus accumbens, and medial preoptic area^10,17,18^. Notably, there are significant sex differences in LBN-induced gene transcription, with few overlapping differentially expressed genes (DEGs) across the sexes. In some cases, these transcriptional sex differences may drive some sex differences in behavior following LBN. For example, adult LBN females show reduced motivation for sucrose, while males show enhanced motivation for sucrose^16^. In other cases, these gene changes may reflect sex differences in molecular mechanisms that converge on the same outcome, such as in the OFC, where LBN affects the glutamate pathway across sexes but via the regulation of different genes in males and females^17^. Thus, it is crucial to consider biological sex when determining the molecular signatures of LBN.

Given the prominent role of the mPFC in mediating behaviors altered by LBN in adulthood and the lasting effects of LBN on gene transcription in other brain regions, here we wanted to determine how LBN altered gene transcription in the adult mPFC of male and female rats. Prior transcriptional studies with this model mainly assessed gene changes in bulk tissue, but here we advanced this work by employing single-nucleus RNA seq (snRNAseq) to identify cell-type-specific transcriptional changes induced by LBN. snRNAseq is particularly advantageous in a heterogeneous region like the mPFC, which has a laminar structure comprised of different classes of excitatory and inhibitory neurons, as well as glial and other cells. This advanced discovery-based approach revealed that, relative to control rats, LBN caused the largest changes in DEGs in vascular cells in the mPFC; with the effect observed in adult female but not male rats. The brain vasculature, which includes endothelial and mural cells such as vascular smooth muscle cells and pericytes^19^, is essential for delivering oxygen and nutrients and supporting neural function. Because the early postnatal period is a critical window for cerebrovascular maturation^20^, these findings raised the possibility that early resource scarcity may alter the developmental trajectory of mPFC vasculature, particularly in females. To test this hypothesis, we used immunohistochemistry and structural analyses to examine vascular organization in the adult mPFC. These studies revealed that LBN reduced vessel coverage, length, and branching selectively in females. Together, these findings identify brain vasculature as a previously unrecognized, sex-biased target of early resource scarcity in the mPFC.

## Methods

### Subjects & Limited Bedding & Nesting Manipulation

Male and female adult Long-Evans rats (Charles River Laboratories, Wilmington, MA or Kingston, NY) were bred in-house to avoid confounding stress from shipping pregnant females. Dams were cohoused for two weeks with male breeders and then removed for the remainder of the pregnancy. All dams were primiparous. Pups were born on PND 0, and on PND 2, litters were culled to 10 pups, five males and five females when possible. Dams and pups were randomly assigned to either control housing or LBN housing in standard shoebox cages with filter tops. Control housing included nesting material, direct access to bedding (Bed-o’Cobs), and one enrichment tube. LBN housing consisted of 1 paper towel for nesting, no enrichment, and a metal grate preventing access to bedding^11^. Rats were in either control or LBN housing through PND9, then put into control cages for the remainder of the study. Rats were weaned at PND 21 and put into sibling co-housing (same-sex and same condition). Adult subjects PND 60-120 were used. All experiments were approved and in accordance with Temple University’s and Georgia State University’s Institutional Animal Care and Use Committee.

#### Single nucleus RNA sequencing

Male and female rats exposed to LBN or control housing were sacrificed at PND 60-120, and brains were flash frozen. Nuclei suspensions derived from fresh frozen mPFC punches were prepared and used single nuclei transcriptomics using the 10x Genomics 3’ gene expression assay (v3.1), according to manufacturer protocols and similar to our prior descriptions^21–24^, and the resulting libraries were sequenced on an Illumina NovaSeq 6000. CellRanger (v3.1.0) was used to deconvolute and align transcriptomics libraries to the reference genome. Raw and filtered expression matrixes were used for quality control and clustering in Seurat as we have described^25–27^. Differential expression was determined using a pseudo-bulk approach via Libra, with a corrected p-value < 0.05 considered significant. Pathway and enrichment analyses were conducted as described.

### Immunohistochemistry

A different cohort of subjects were used for morphological analysis. Adult rats were anesthetized and perfused with ice-cold PBS followed by 4% paraformaldehyde (PFA). Brains were post-fixed for 24h in 4% (PFA) and transferred into 30% sucrose at 4°C for 48 hours and cyrosectioned coronally at 35 μm on a Leica CM3050 cryostat (Leica Biosystems). Brain sections were collected and stored in cryoprotectant (30% Sucrose and 30% glycerol in Phosphate Buffered Saline, PBS) at 4ºC until further use. Brain slices (free-floating) were incubated with blocking solution (10% Donkey serum, 0.4% Triton in 1X PBS) for 1h at room temperature (RT). Then, brain sections were incubated with primary endothelial cell marker, CD31(R&D systems; #AF3628, 1:250), in 1% Donkey-PBS-T at 4 °C for 48 h under agitation. After washing, sections were incubated with a secondary antibody diluted in 1% Donkey-PBS-T 0.4% (Jackson Immuno, 1:250) conjugated to Alexa Fluor 488 for 3h at 4 ºC with agitation, and nuclei were stained with ProLong™ Gold Antifade Mountant (ThermoFisher, P36930) After PBS washes, sections were mounted using anti-frost slides.

### Confocal imaging and image analysis

Confocal images were acquired using a Zeiss LSM 980 microscope (Carl Zeiss, Germany) with ZEN LTE software (v3.11 Carly Zeiss, Germany) and LSM Plus processing. mPFC images were taken in both infralimbic and prelimbic regions of both hemispheres. Images were taken using 20× objective with a z-step size of 0.4 μm. Images then underwent post-processing background subtraction in ZEN LTE software with a 100 μm rolling ball radius for all images. For morphological evaluation, we used IMARIS software (version 10.2.0 Bitplane, Belfast, UK). Briefly, volume and area were quantified by applying 3D surface rendering, while vessel length and branch points were measured using filament tracing of the confocal image stacks in their respective channels. Identical settings, including fixed intensity thresholds and voxel sizes, were applied within each experiment to ensure consistency.

### Statistical Analysis

Statistical analyses were performed using GraphPad Prism (GraphPad Software, San Diego, CA, USA) and Python (v. 3.12.12; Python Software Foundation). Morphological measures were analyzed using two-way analysis of variance (ANOVA) with sex and condition (LBN vs. control) as between-subjects factors. Based on our snRNAseq findings, we *a priori* hypothesized that LBN would alter vascular morphology in females; therefore, planned comparisons were conducted to test for condition effects within each sex. Alpha was set at *p* < .05.

## Results

### Early resource scarcity broadly increases differently expressed genes in females

To characterize early-life stress and sex-specific transcriptomic changes across cell types in the mPFC, we utilized snRNAseq to generate nuclear transcriptome profiles, and we identified cell types expected for the region (Fig 1A and 1B). DEGs were identified by comparing adult male and female rats exposed to LBN conditions to male and female rats to male and female control rats raised in a normal housing environment, in a sex aggregated (Fig 1C left and Table S1) and sex disaggregated fashion (Male: Fig 1C middle and Table S2; Female: Fig 1C right and Table S3). LBN females generally had increased DEGs across most cell types compared to LBN males. In the vascular cell type, females had 46 DEGs, the highest of any cell type, whereas males had no differential expression in this cell type. Females also showed differential expression in intratelencephalic neurons across the cortical layers, with the altered expression profile of each layer being largely unique (Fig 1D).

**Figure 1.**
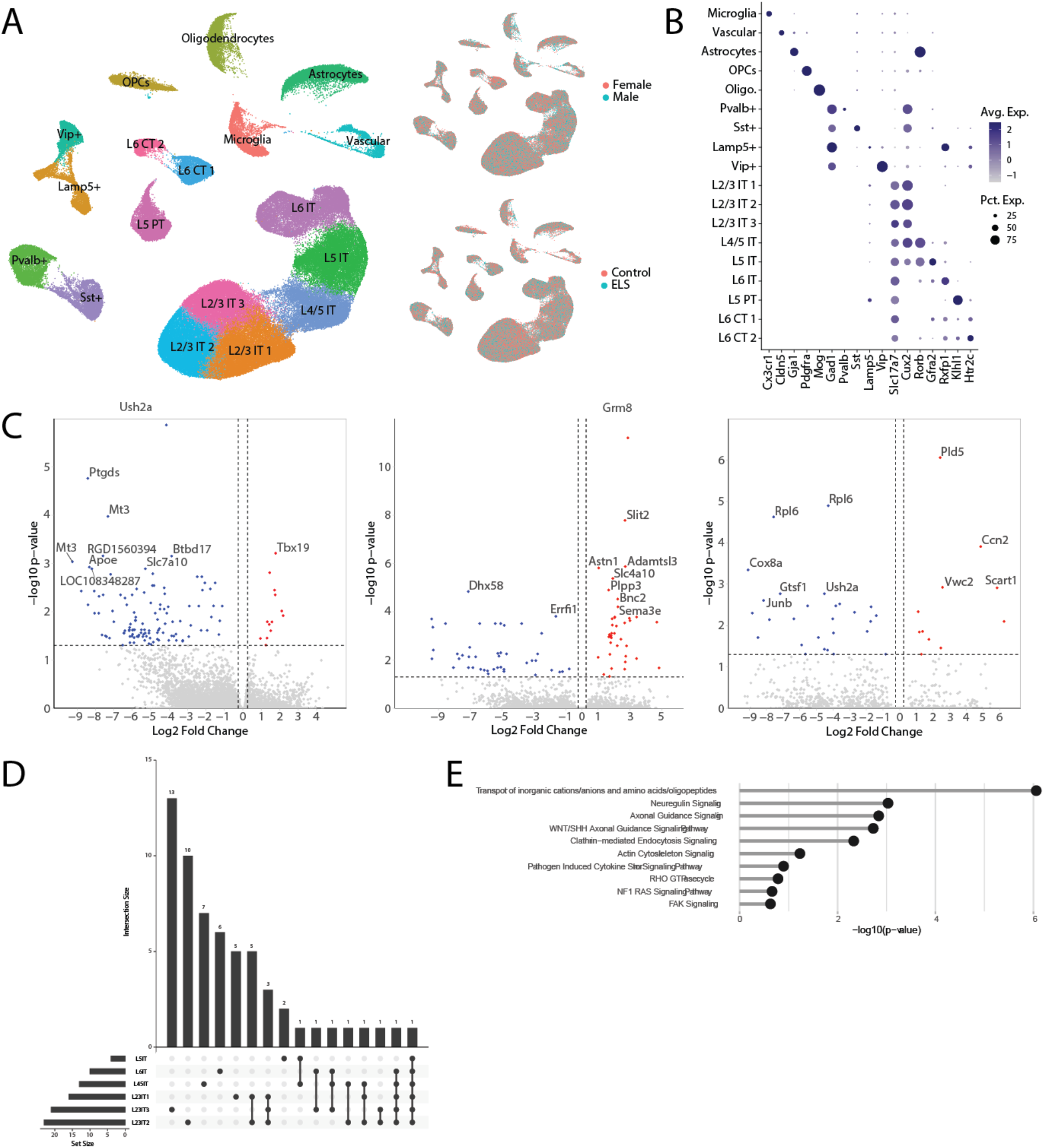
Single nuclei transcriptomics of the mPFC from male and female rats exposed to ELA via the LBN model. **A)** A uniform manifold approximation and projection of the cellular cluster from the mPFC, with no difference in representation seen by treatment or sex. **B)** A dot plot showing example marker genes used to identify mPFC cell types. **C)** Volcano plots showing differentially expressed genes from all clusters for the sex aggregated (left), male (middle), and female (right) analyses. **D)** LBN-induced alterations of transcriptional programs in intratelencephalic neurons were largely cortical layer-specific. **E)** Example altered canonical signaling pathways in nuclei from the vascular cluster in females.

### LBN alters the genetic profile of female vascular cells in the mPFC

Our snRNA-seq dataset revealed that our vascular cell cluster in the mPFC was influenced by LBN only in females. This vascular cluster was identified by CLDN5 expression, which means it is largely composed of endothelial cells, but could include vascular smooth muscle cells and pericytes. To identify the molecular pathways that might be altered, we ran a pathway analysis of our identified DEGs using Ingenuity Pathway Analysis (IPA; Fig 1E and Table S4). Here we found pathways important for blood-brain barrier transport, (Transport of inorganic cations/anions and amino acids/oligopeptides, Cathrin-mediated Endocytosis Signaling), angiogenesis and endothelial morphology, (Neuregulin Signaling, Axonal Guidance Signaling, WNT/SHH Axonal Guidance Signaling Pathway, Actin Cytoskeleton Signaling, RHO GTPase cycle) and immune signaling (Pathogen Induced Cytokine Storm Signaling Pathway, NF1 RAS Signaling Pathway, FAK Signaling) were altered. *In silico* modeling of regulatory networks controlling the altered transcriptional program observed in the vascular cluster identified 4 putative upstream transcriptional regulators, *Foxc1, Pax6, Arrb1*, And *Zfp36* as potential direct mediators of stress-specific changes in endothelial structure. Collectively, these transcriptional signatures pointed to disruption of endothelial cytoskeletal architecture, vessel patterning, and barrier-related transport functions, predicting that LBN exposure would result in measurable alterations in cerebrovascular morphology. This provided a strong rationale for direct structural analysis of brain vasculature.

### LBN reduces vasculature coverage and complexity in the mPFC of females but not males

To test the prediction from our snRNA-seq analyses that LBN exposure would alter endothelial structural organization, we next examined cerebrovascular morphology in the mPFC (Fig 2A). Because the early postnatal window coincides with rapid maturation of the brain vasculature and undergoes active vascular remodeling^20^, we hypothesized that disruption during this period would result in lasting changes to endothelial network architecture. To investigate how LBN affects vascular structure in the mPFC, we utilized 3D reconstruction of immunohistochemistry staining for endothelial cells (Fig 2B). There were no differences in the number of endothelial vessels across all groups (LBN: *F*_1,20_ = 1.235, *p* = .280; Sex: *F*_1, 20_ = 4.55, *p* = .045; interaction: *F*_1,20_ = 0.5767, *p =* .457) (Fig 2C). However, volumetric analysis revealed that LBN decreased vessel coverage of the mPFC in females only (LBN: *F*_1, 20_ = 3.00, *p* = .098; interaction: *F*_1,20_ = 3.31, *p* = .084, Planned comparisons *p* = .027) (Fig 2D). Mean volume per was also decreased in females (LBN: *F*_1, 20_ = 2.454, *p* = .1329; interaction: *F*_1,20_ = 6.121, *p* = .0224, Planned comparisons *p* = .0125)(Fig 2E) The rodent mPFC has two main sub-regions, the infralimbic (IL) and prelimbic (PL) cortices. We ran a three way ANOVA and found no region effects but LBN (F_(1, 40)_ = 4.660, *p* = .0379) and Sex × LBN (F_(1, 40)_ = 5.43, *p* = .025) effects leading us to separate by sex and run 2×2 ANOVA in which we identified a main effect of LBN in females but not males across these regions (Female (Region × Condition); LBN: F_(1, 18)_ = 6.07, *p* = .024) (Suppl. Figure 1). Given that we did not identify sub-region-specific effects of LBN, the remaining analysis pooled images from the IL and PL.

**Figure 2.**
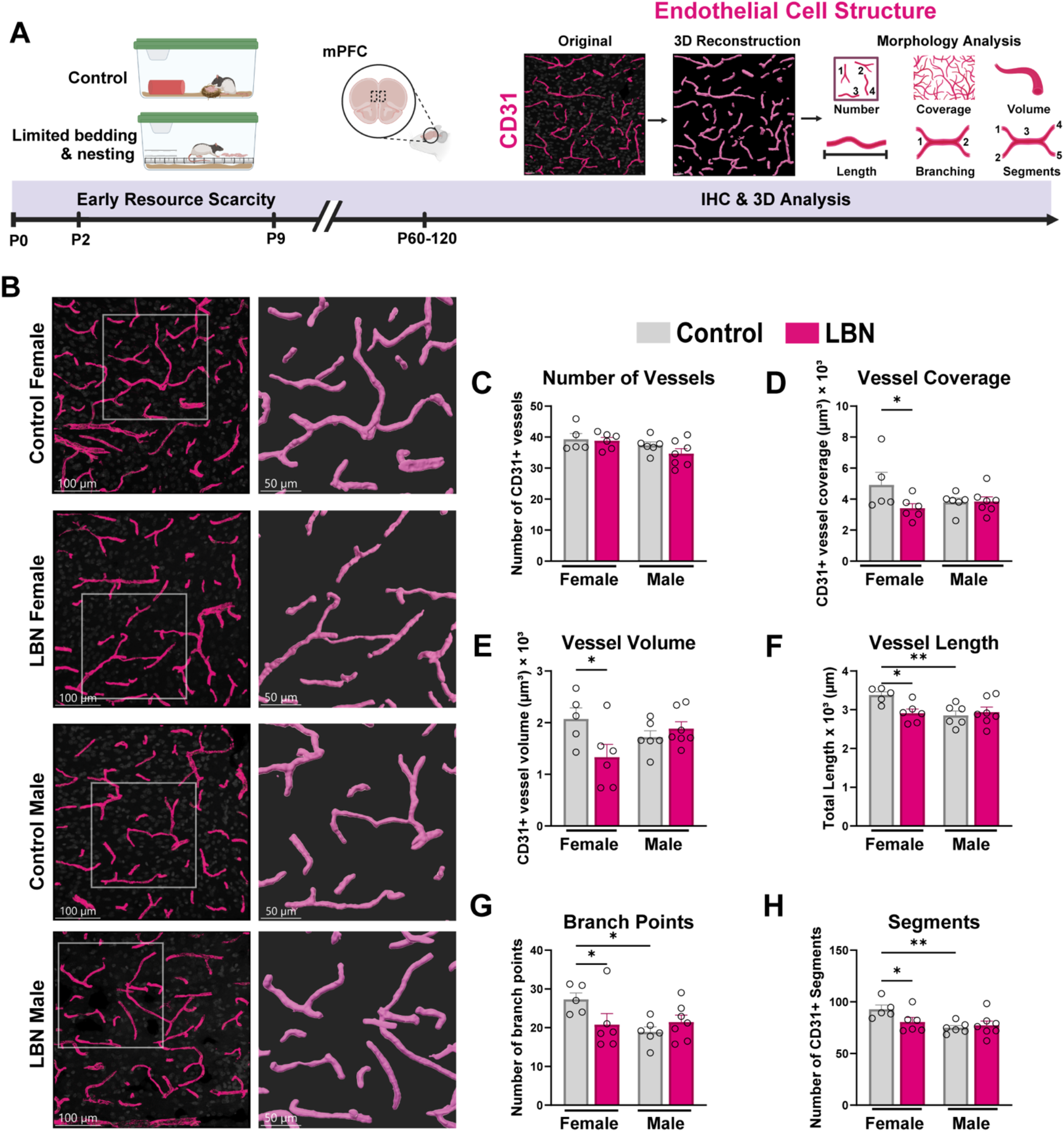
Early-life resource scarcity reduces vascular coverage and network complexity in the adult female mPFC. **A)** Experimental timeline of limited bedding and nesting (LBN) or control conditions from postnatal day (P)2–9. Animals were aged to adulthood, after which medial prefrontal cortex (mPFC) tissue was collected and immunostained for the endothelial marker CD31. Confocal image stacks were subjected to Imaris-based 3D reconstruction for quantitative vascular morphometric analysis. **B)** Representative 20× confocal images of CD31 immunolabeling in the mPFC from control and LBN-exposed females and males (Scale bar 100 μm) with corresponding Imaris-generated 3D reconstructions (80% zoom, Scale bar, 50 μm). **C)** Total number of CD31^+^ vessels. **D)** CD31^+^ vascular coverage. **E)** Mean volume of CD31^+^ vessels. **F)** Mean length of CD31^+^ vessels. **G)** Mean number of branch points. **H)** Mean number of vascular segments. Graphs display mean ± SEM with individual animals represented as data points (n = 5–7 per group, 4-12 images/animal). Statistical analysis was performed using two-way ANOVA followed by planned comparisons (#*p* < .07, \**p* < .05, \*\**p* < .01).

To identify the structural features that contribute to decreased coverage, we used filament analysis to identify changes in vessel length, branching, and diameter. LBN decreased vessel length in females and demonstrated baseline sex differences (LBN: F_(1, 20)_ = 2.81, *p* = .109; Sex: F_(1, 20)_ = 4.61, *p* = .044; Interaction: F_(1, 20)_ = 5.676, *p* = .0272. Post hoc, Female: Control vs LBN *p* = .0122; Control: Female vs Male *p* = .006)(Fig 2F). We observed decreased branching in LBN females and a baseline sex difference in branching by number of branch points (Sex: p = .0715; Interaction: F(1, 20) = 4.99, *p* = .037. Post hoc, Female: Control vs LBN *p* = .042; Control: Female vs Male *p* = .011)(Fig 2G) and by the total number of segment (Sex: F(1, 20) = 7.42, *p* = .013; Interaction: F(1, 20) = 3.54, *p* = .075. Post hoc, Female: Control vs LBN *p* = .0488; Control: Female vs Male *p* = .0062)(Fig 2H). Our identified baseline sex difference with control females having longer and more complex vessels than males aligns with prior reports of greater baseline brain capillary networks in females than males^28^. There are no significant differences in mean capillary diameter across groups (Suppl. Figure 1A).

## Discussion

Prior work has demonstrated that LBN disrupts cognition and motivated behaviors mediated by the mPFC, often in a sex-biased manner^10,16,29,30^. However, the cellular mechanisms underlying these persistent effects remain poorly understood. Here, we identify, for the first time, cell-type-specific transcriptional changes in the adult mPFC following postnatal resource scarcity in males and females. LBN induced more DEGs across pyramidal neuron subtypes of the mPFC in females than in males. However, the most pronounced transcriptional changes occurred in vascular cells, and these effects were observed exclusively in females. This finding led us to test whether postnatal resource scarcity alters vessels in the mPFC of females. We replicated some baseline sex differences in mPFC vessel morphology. However, our main finding was that LBN selectively reduced vessel coverage, length, and structural complexity in females, indicating persistent disruption of the mPFC capillary network. While most prior work has investigated how changes in the early environment alter neurons, our findings highlight brain vasculature as an additional and previously underappreciated target of changes in the postnatal environment. The early postnatal period is a critical window of cerebrovascular growth and vascular maturation^20,31^ . Our data reveal that early resource scarcity produces persistent, female-specific disruption of mPFC vasculature, identifying a novel pathway through which early adversity may reprogram the brain to affect later brain function and disease vulnerability.

### LBN altered gene transcription intratelencephalic (IT) neurons of the female mPFC

The mPFC exerts top-down control over downstream targets via two main types of output neurons: pyramidal tract (PT) and intratelencephalic (IT) neurons^32–34^. Modest LBN-induced gene changes were observed in PT neurons across sexes. In contrast, LBN produced pronounced transcriptional alterations in IT neurons in females, within layers 2/3, 4/5, and 6. Here, we molecularly defined three subclusters of L2/3 IT neurons, and in all three clusters *C1qb*, which encodes a subunit of the complement protein C1q, was downregulated in LBN v. control females. During development, C1q is a crucial signal for synaptic pruning by microglia^35,36^. In other regions, such as the paraventricular nucleus of the hypothalamus, LBN impairs microglial-mediated synaptic pruning, resulting in aberrant glutamatergic connectivity^37^. Thus, the downregulation of *C1qb* in L2/3 IT neurons may indicate that similar disruptions in microglial-synaptic interactions may occur within the mPFC following early resource scarcity. Beyond its role in synapse regulation, C1q also modulates neuroimmune function by limiting inflammation and promoting a shift of microglia and macrophages toward an anti-inflammatory response^38,39^. Another neuroimmune gene^40^, *Dhx58*, was also downregulated in LBN females across all IT layers, further supporting altered immune signaling to these neurons. One upstream regulator of these and other DEGs in L2/3 IT neurons is Tripartite motif-containing protein 32 (TRIM32). TRIM32 promotes anxiety and depressive behavior in adult mice exposed to chronic mild stress^41^. Together, these findings suggest that TRIM32 may represent a molecular link between inflammatory and behavioral outcomes of early-life adversity in females. Given that IT neurons project extensively to other cortical regions, as well as to the amygdala and striatum^32–34^, LBN-induced transcriptional alterations in these neurons may contribute to changes in cognition, affect, and motivated behaviors, particularly in females.

### LBN altered genes in female mPFC vasculature involved in angiogenesis and immune function

Our most robust LBN-induced gene changes were observed only in our female vascular cells. IPA identified many signaling pathways, most notably those related to angiogenesis and immune signaling. Within the angiogenic pathways, we identified a subset of upregulated genes, *Slit2, Sema3e*, and *Itgb8*, typically associated with axon guidance but also exert potent regulatory effects on angiogenesis^42–44^. A recent study identified the repulsive axon guidance cue *Slit2* as a potential “braking” mechanism that inhibits angiogenesis by reducing vessel overgrowth^31^. Here, we find that LBN increased the expression of *Slit2* only in females, suggesting enhanced signaling that may constrain vascular growth. *Sema3e* has previously been identified as anti-angiogenic^45^ and is also increased in LBN vs. control females. Recent work further implicates *Sema3e* in postnatal vascular refinement and regional specialization^31^, consistent with a role in regulating vascular architecture. *Itgb8* encodes the β8 subunit of the αvβ8 integrin; αvβ8 is required for normal brain vascular morphogenesis and restricts excessive endothelial sprouting^46^. Pathway analysis also links *Itgb8* to focal adhesion kinase (FAK) and actin cytoskeleton signaling. Integrin controls endothelial stability through activation of FAK and consequent suppression of RhoA-mediated cytoskeletal tension^47^. Consistent with this, Rho GTPase signaling, central to endothelial branching and cytoskeletal tension, is also altered in LBN vascular cells. RhoA is critical for vessel formation, regulating angiogenesis and vascular development^48^. Collectively, the upregulation of *Slit2, Sema3e*, and Itgb8, together with alterations in Rho GTPase signaling, indicates coordinated LBN-induced modulation of pathways governing endothelial sprouting and vascular structure in females. These molecular changes align with the reduced vessel length and branching observed in the mPFC of adult LBN females.

LBN vascular nuclei also demonstrate enrichment of pathogen-induced cytokine storm signaling and phagosome formation, pathways similarly associated with stress-related inflammation in humans^49,50^ and in early-life stress models in rats^51^ . Thus, LBN may elicit an inflammatory endothelial response. In fact, *Kit* was downregulated in the vasculature of LBN females, suggesting a loss of an endothelial anti-inflammatory regulator known to limit NF-κB-driven cytokine amplification^52^, which may sensitize CNS vessels to inflammatory cues. Together, these DEGs suggest that LBN induces a female vascular transcriptional profile consistent with heightened inflammatory signaling and increased endothelial sensitivity to immune challenge.

We identified several upstream transcriptional regulators, including *Arrb1, Foxc1, Pax6*, and *Zfp36*, that may contribute to the observed LBN-associated vascular gene expression changes in females. Conditional endothelial-specific knockout of *Arrb1* results in reduced capillary branching^53^, *Foxc1* is important in the stability of pericytes and endothelial cells during development^54^. *Pax6* is important for astrocyte maturation and increases expression in reactive astrocytes^55^ potentially implicating astrocytes as a mediator for endothelial dysfunction. *Zfp36* is an RNA-binding protein that influences the stabilization of inflammatory mRNAs^56^ and has upstream roles in endothelial cell proliferation and endothelial tip extension during development^57^. Future work will investigate how these transcriptional regulators interact to coordinate endothelial and glial gene programs underlying LBN-associated alterations in angiogenic signaling and vascular maturation.

### LBN reduces vessel coverage and complexity in mPFC of females

Our snRNAseq data indicated potential alterations in mPFC vasculature in adult females, but not males, following LBN. We therefore assessed whether LBN altered mPFC vessel morphology. LBN reduced vascular coverage in the female mPFC, driven by decreases in vessel length and branching. These effects were not subregion-specific and were absent in males. Under control conditions, females exhibited more complex vascular architecture than males, consistent with prior reports demonstrating that females have larger and more highly branched capillary networks than males across multiple brain regions at baseline^58^. Other manipulations decrease vessel coverage and complexity in males^59^, indicating that male vasculature retains the capacity for structural remodeling. Therefore, it is unlikely that control males were at a floor for vessel complexity, but rather that LBN does not alter mPFC vasculature in young adult males^60^. On some measures, LBN reduced female vascular architecture to levels comparable to those of control males. However, equivalence in vessel coverage does not imply equivalence in biological impact. Rather, these findings suggest that LBN shifts the female mPFC away from its normative, sex-specific vascular phenotype. Given that reduced vascular density constrains oxygen and glucose delivery, reducing females to a “male-typical” level likely represents a loss of metabolic capacity for the female mPFC, even if that same level is sufficient in males. Thus, LBN-induced vascular reductions in females may limit energetic support to the mPFC and compromise its ability to sustain optimal function.^61–63^

Other studies on stress effects on vessel morphology are fairly limited^61,64^. A recent study using a chronic unpredictable stress manipulation in adult mice found this stressor also reduced vessel coverage in the mPFC of females, but surprisingly increased it in males^61^. Another early stress manipulation, maternal separation, reduces vessel diameter in the mPFC of adult mice, but other morphology measures were not assessed, and data were not disambiguated by sex^64^. There is more research on adult chronic stress effects on blood-brain barrier (BBB) permeability^61,62,65–68^, where again the female mPFC seems particularly vulnerable to stress-induced disruption^65^. Future studies will determine how LBN affects BBB permeability. Collectively, this research suggests a female-biased vulnerability of mPFC vasculature to stress. The early life manipulations occur before puberty, so perhaps it is the organizational effects of the perinatal surge of testosterone (which occurs before LBN and maternal separation are implemented) that masculinizes the mPFC, increasing resilience to stress-induced vascular changes in males.

### Stress-induced changes in neurovasculature: a female-biased risk factor for brain health

Reduced vascular coverage in the mPFC may constrain oxygen and glucose delivery, limiting metabolic support for executive control processes and increasing vulnerability to brain dysfunction. Vascular integrity is increasingly recognized as a critical determinant of neuropsychiatric and neurodegenerative disease risk^69^.

Cocaine, a potent vasoconstrictor, disrupts cerebral blood flow, impairs endothelial function, and increases the risk for ischemic and hemorrhagic stroke^70,71^. Early vascular remodeling combined with cocaine-induced vascular stress may amplify susceptibility to cerebrovascular injury and small vessel pathology^72^. Although females exhibit delayed vascular aging under typical conditions^73^, emerging clinical and epidemiological evidence indicates that early-life adversity may disproportionately accelerate vascular aging in females^46^. Clinical studies linking early stress to female-biased cerebrovascular disruption remain limited; however, peripheral vascular dysfunction is more severe following early-life trauma in women ^74^. Together, these findings position early-life stress as a female-biased risk factor for later-life cerebrovascular vulnerability and underscore the need for translational studies examining its impact on vascular aging and cerebrovascular disease trajectories.

The postnatal period represents a developmental window in which much of neuronal proliferation and migration has stabilized, yet the brain vasculature continues to undergo active growth and remodeling^20,75^. Despite this dynamic vascular maturation, the impact of the early environment on cerebrovascular development remains understudied. Here, we demonstrate persistent, female-specific vascular remodeling in the mPFC following early resource scarcity. These findings broaden the conceptual framework of early-life risk beyond neuron-centric models and identify the cerebral vasculature as a sex-specific substrate through which early adversity may program long-term vulnerability to psychiatric and cerebrovascular disease.

## Acknowledgements

This work was supported by the National Institutes of Health (R21 DA059966, R01 DA049837, R01 DA056534, R34 DA061483, R21 DA062844, S10OD032336) and the National Science Foundation (IOS-2313253). The content is solely the responsibility of the authors and does not necessarily represent the official views of the NIH.

