## Supplemental Figures for "Early resource scarcity drives persistent transcriptional changes and vascular remodeling in the female prefrontal cortex"

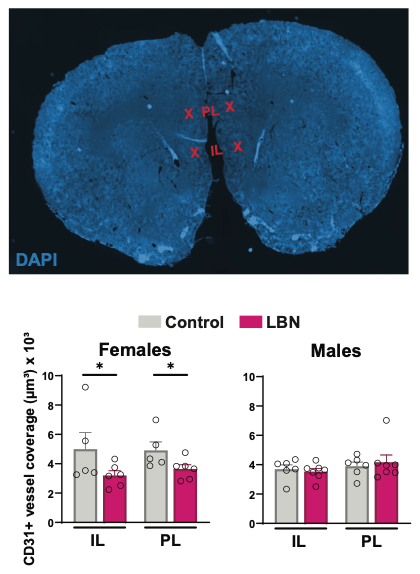
**Supplemental Figure 1: CD31 coverage in subregions of the mPFC**

A) Representative image 5× Tile scan image of DAPI signal demonstrating subregion images Infralimbic (IL) and prelimbic (PL). B-C) Three-way ANOVA was used to identify condition × sex × region significance in CD31 coverage (n=5-7, 4-12 images/animal). Three way ANOVA result revealed main effect of Condition (*p* = .03) and Sex × Condition (*p* = .02). Data was then split by sex and 2-way ANOVAs were run to compare IL vs PL control and LBN. Bars represent mean ± S.E.M. Each individual animal is represented as data point (*p* < .07, **p* < .05, ***p* < .01) B) Female Two-way ANOVA revealed main effect of LBN on CD31 vessel coverage (*p* = .02). C) Male Two-way ANOVA showed no significant effect of CD31 vessel coverage

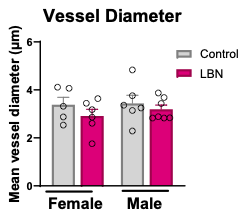

**A**

**Supplemental Figure 2: CD31 Mean Diameter**

A) Mean diameter of CD31+ segments. Graphs display mean ± SEM with individual animals represented as data points (n = 5–7 per group, 4-12 images/animal). Statistical analysis was performed using two-way ANOVA followed by planned comparisons.
